# RNA structure prediction including pseudoknots through direct enumeration of states

**DOI:** 10.1101/338921

**Authors:** Ofer Kimchi, Tristan Cragnolini, Michael P. Brenner, Lucy J. Colwell

## Abstract

The accurate prediction of RNA secondary structure from primary sequence has had enormous impact on research from the past forty years. While many algorithms are available to make these predictions, the inclusion of non-nested loops, termed pseudoknots, still poses challenges. Here, we describe a new method to compute the entire free energy landscape of secondary structures of RNA resulting from a primary RNA sequence, by combining a polymer physics model for the entropy of pseudoknots with exhaustive enumeration of the set of possible structures. Our polymer physics model can address arbitrarily complex pseudoknots and has only two free loop entropy parameters that correspond to concrete physical quantities, over an order of magnitude fewer than even the sparsest state-of-the-art algorithms. Our model outperforms previously published methods in predicting pseudoknots, while performing on par with current methods in the prediction of non-pseudoknotted structures. For RNA sequences of ~ 45 nucleotides, or ~ 90 with minimal heuristics, the complet–e enumeration of possible secondary structures can be accomplished quickly despite the NP-complete nature of the problem.

---

RNA molecules play physiological roles that extend far beyond translation. In human cells, most RNA molecules are not translated [1]. Non-coding RNAs interact functionally with mRNA [2], DNA [3], and proteins [4], and can be as large as > 200 nucleotides (ntds) [5, 6]. However, a substantial fraction are < 40 ntds in length, including miRNAs and siRNAs, which serve as regulators for the translation of mRNA [2, 7], and piRNAs which form RNA-protein complexes to regulate the germlines of mammals [8]. The *in vitro* evolution of RNA, especially through SELEX [9–11], has led to an explosion of applications for short RNA molecules, due their ability to tightly and specifically bind to a remarkable range of target ligands [12].

Overwhelmingly, the properties of short non-coding RNA molecules are tied to their three-dimensional, or tertiary, structures [5, 13–16]. Such structures are formed because of the energetic favorability of bonds between complementary nucleotides (primarily A to U, C to G, and G to U). However, these bonds impose an entropic cost; therefore, the conformations most frequently adopted balance the energetic gain of maximal base-pairing with the entropic cost of structural constraints. In equilibrium, the RNA adopts each possible structure with Boltzmann weighted probabilities.

Because of the relevance of RNA structure to function [17, 18], current research aims to predict the minimum free energy structures given the sequence. Algorithms typically predict “secondary structure”, a list of the base pairings [19]. The early Pipas-McMahon RNA structure prediction algorithm sought to completely enumerate and evaluate the free energy of all possible secondary structures, thereby constructing the entire energy landscape [20]. This NP-complete approach was quickly supplanted by dynamic programming, which has since dominated RNA structure prediction [21–25]. These algorithms efficiently consider an enormous number of structures without explicitly generating them, by iteratively finding the optimal structure for subsequences [26].

However, such algorithms have difficulty predicting RNA secondary structures that include pseudoknots, i.e. structural elements with at least two non-nested base pairs (see Fig. S1A for an example) that make up roughly 1.4% of base pairs [26] and are overrepresented in functionally important regions [27] of RNA. Pseudoknots are disallowed from the most popular RNA structure prediction algorithms (e.g. Refs. [28–30]) due to computational cost; indeed, structural prediction including all pseudoknots has been shown to be NP-complete [31–33]. Significant advances have been made with heuristics, which do not guarantee finding the minimum free energy structure [34–38], and by disallowing all but a narrow class of pseudoknots [39–46].

**FIG. 1:**
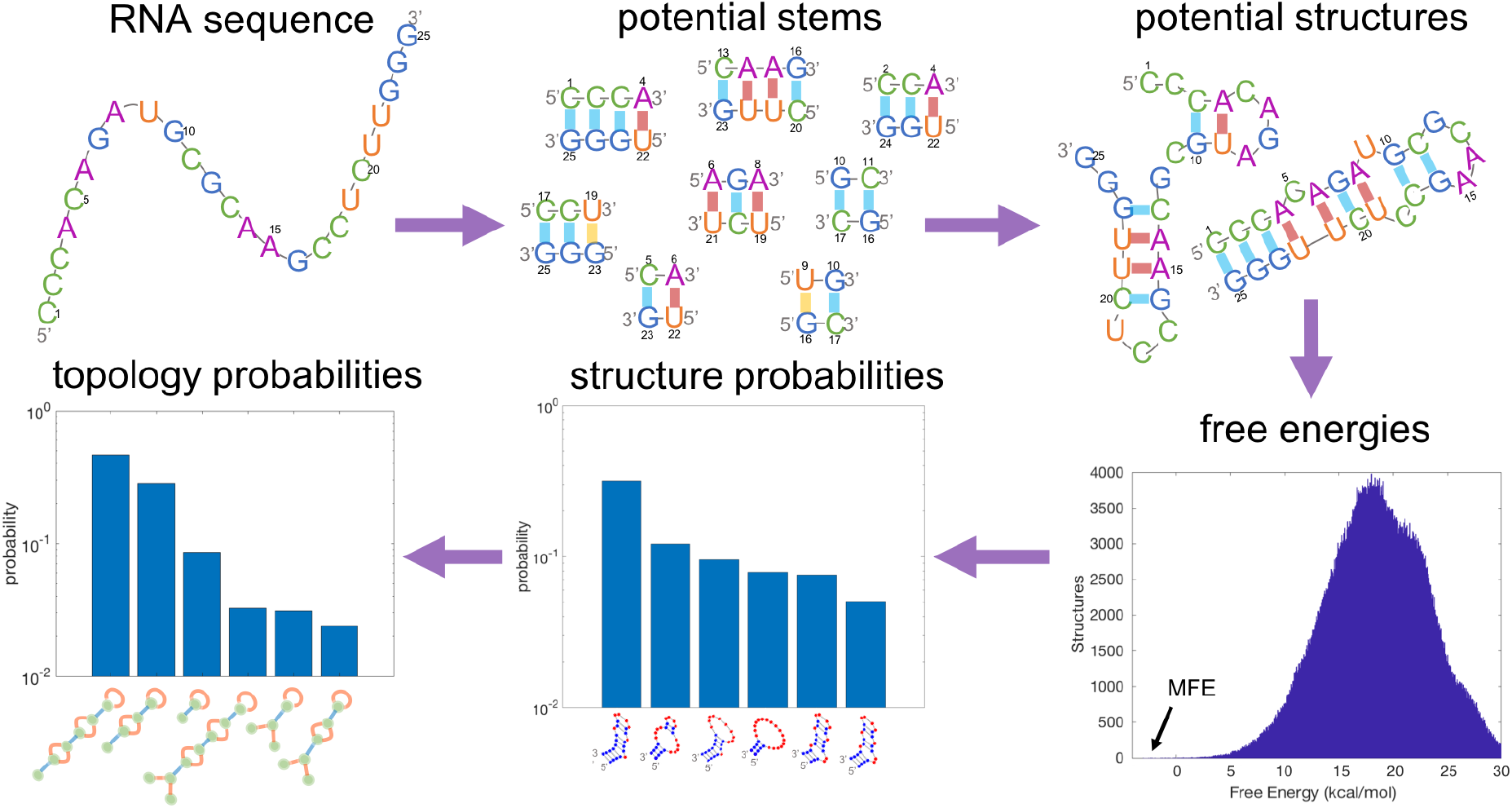
Schematic overview of the algorithm. Given an RNA sequence, the algorithm first enumerates all potential stems (sequences of base pairs) which can form. It then searches for all possible combinations of stems, such that no nucleotide is paired with more than one other, thus forming all possible secondary structures. For each structure, it calculates the free energy, which is comprised of a bond energy term and an entropy term. The histogram of free energies for the sequence shown is plotted with an arrow pointing to the Minimum Free Energy (MFE). Given the entire free energy landscape, the algorithm calculates the probability of any arbitrary secondary structure of forming in equilibrium. Finally, we coarse grain over similar structures described by the same topology (described in Section III), arriving at a probability distribution for every possible topology forming in equilibrium.

A major challenge for predicting pseudoknotted structures is the relative lack of experimental data [47]. Thus, up until recently, theoretical approaches have largely been limited to simple H-type pseudoknots [39, 45, 48, 49]. A recent strategy uses machine-learning of large experimental datasets [45, 50, 51]. Although these approaches can be useful, they come with the disadvantages of compounding possible experimental errors, and often using an enormous number of parameters which can hamper generalizability. A sketch of a theoretical description of pseudoknot entropies based on polymer physics was developed by Isambert and Siggia [34, 52]; however, their derivations have not been published.

In this study, we demonstrate that for short RNA sequences, it is possible to exactly solve for the probability that the RNA will fold into any given structure, in eluding those with pseudoknots. Complete enumeration of the RNA structure landscape is feasible even for biologically relevant RNA sequences (Section I). Our approach combines a method based on the work of Isambert and Siggia (Section II) with a novel graph-theoretical depiction of the RNA (Section III) to exactly calculate the entropy of each structure, treating both pseudoknotted and non-pseudoknotted RNA structures equivalently. The entropies of structures of arbitrary complexity can be analytically computed with just two experimentally derived physical parameters: the persistence length of single-stranded RNA, and the volume within which two RNA nucelotides are considered bound. This represents an enormous parameter reduction compared to state-of-the-art algorithms like the Cao-Chen or Dirks-Pierce models, which have 258 and 11 parameters, respectively, for H-type pseudoknots alone, and ~ 18 parameters for non-pseudoknotted loops [51]. We test our model predictions on molecules from the RNAStrand [53], PseudoBase++ [54], and CompraRNA [55] databases and find good agreement with experimental results (Section IV). Although we fit our entropy model to data from non-pseudoknotted structures, we find that our model outper-forms previously published methods in predicting pseudoknots, while performing on par with current methods in the prediction of non-pseudoknotted structures.

## I. ENUMERATING RNA STRUCTURES

The Pipas-McMahon algorithm [20] first enumerates all possible secondary structures for a given sequence (*sans* pseudoknots), and then evaluates the free energy for each, to construct the entire free energy landscape for non-pseudoknotted structures. A major shortcoming is the significant computer time required for long sequences. However, the exponential increase in computer power over the past forty years, coupled with increased appreciation for the physiological and engineering relevance of short RNA strands suggest revisiting this approach. In this section, we describe the process by which we exhaustively enumerate the secondary structures into which an arbitrary given sequence can fold. We first number the nucleotide sequence from 1 to *N* from the 5’ end. We define an *N* × *N* symmetric matrix *B* which describes which nucleotides can bind to each other: *B_i,j_* = 1 if nucleotides i and j can bind to make base pair *i · j* (i.e. they belong to the set {(*A,U*), (*C,G*), (*G,U*)}), and 0 otherwise.

Next, we search for all possible stems (strings of consecutive base pairs) that could form. We define a parameter m to be the minimum allowed stem length (*m* ≥ 1; *m* = 1 throughout unless otherwise specified). We also impose the physical constraint that hairpins (single-stranded region connecting one end of a stem) have a minimum length of 3 nucleotides. We include not only the longest possible stems that can form, but all contiguous subsets of those stems [56, 57]. We denote the number of stems found by *N*_stems_.

We next define the *N*_stems_ × *N*_stems_ symmetric compatibility matrix *C*, where *C_p,q_* = 1 if a structure could be made with both stems *p* and *q* (*C_q,q_* = 1 ∀ *q*). We impose the constraint that each nucleotide may be paired with, at most, one other nucleotide by setting *C_p,q_* = 0 if stems p and q share at least one nucleotide.

Finally, we explicitly enumerate the remaining possible secondary structures by identifying all compatible combinations of stems. Starting from a single stem *s*_1_, we consider stems *s*_2_ where 1 ≤ *s*_1_ < *s*_2_ ≤ *N*_stems_ and add the first stem for which *C*_*s*_1_,*s*_2__ = 1. Then, we repeat the process, adding the first stem *s*_3_ > *s*_2_ compatible with both *s*_1_ and *s*_2_, and so forth, continuing until we can add no more stems. We add the resulting structure, composed of say *M* stems, to the list of possible structures, then remove the last stem added (to obtain the structure composed of stems *s*_1_, *s*_2_, …, *s*_*M*−1_) and continue the process. This algorithm returns all possible secondary structures resulting from the primary sequence.

The algorithm described here was implemented in MatLab and all code is available on the GitHub repository https://github.com/ofer-kimchi/RNA-FE-Landscape.

Having completely enumerated the possible secondary structures, we calculate the probabilities that the RNA will fold into each of them by calculating their free energies.

## II. CALCULATING FREE ENERGIES

The probability of the RNA sequence folding into a given equilibrium structure *σ* is given by the Boltzmann factor

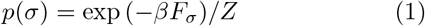

where *β* = 1/*k_B_T* (*T* is the temperature and *k_B_* is Boltzmann’s constant), and the partition function, *Z*, is defined such that the probability distribution is normalized: Σ_*σ*_ *p*(*σ*) = 1. Here *F_σ_*, the free energy of structure *σ*, is a function of the energy *E_σ_* and entropy *S_σ_* of the structure:

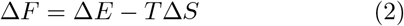

where we drop the subscripts for notational convenience and introduce Δs to signify that free energies are measured with respect to the free chain. We separate the free energy calculation into the free energy of stems and the free energy of loops.

### A. Calculating bond energies

We make the simplifying assumption that the energy Δ*E* in Eq. (2) is determined solely by the base pairs in the structure, ignoring higher order corrections to the energy. Thus, each stem, *s*, contributes an energy Δ*E_S_* such that Δ*E* = Σ_*s*_ Δ*E_S_*. To calculate the terms ΔE*S*, we consider nearest-neighbor interactions among base pairs [58]. Previous work has shown it reasonable to include (whenever appropriate) the contribution of unpaired nucleotides on both sides of each stem in the nearest-neighbor terms for the first and last base pairs of the stem [25]. Specifically, we used tabulated parameters for Δ*H* from Refs. [50, 59, 60], well documented by Turner and Mathews in the Nearest Neighbor Database [61]. Our entropy model (described below) was used in place of the entropies of hairpin, bulge, internal, and multibranch loops and we set the enthalpy terms of these loops (aside from nearest-neighbor interactions) to zero; we did not consider mismatch-mediated coaxial stacking, symmetry penalties or penalties for specific closures of stems; and we implemented coaxial stacking terms in place of terminal mismatches or dangling ends whenever possible in multibranch loops.

### B. Calculating entropies

Entropies are calculated as being comprised of two independent parts: the entropic cost of forming stems and the entropic cost of forming loops, such that Δ*S* = Δ*S*_loops_ + Σ_stems_ Δ*S*_stem_.

The entropies of stems represent the entropy lost when an RNA forms base pairs. This entropy is considered in the same fashion as the energetic parameters (each energetic parameter has an accompanying entropic parameter). Therefore, as for the energies, the entropic parameters consider pairwise RNA base pair interactions, and Δ*S*_stem_ thus depends on the specific nucleotides comprising the stem. In contrast, we make the approximation that Δ*S*_loops_ is independent of the identities of the nucleotides comprising the single-stranded regions.

## III. CALCULATING LOOP ENTROPIES: RNA FEYNMAN DIAGRAMS

We model single-stranded regions comprised of x unpaired nucleotides (ntds) as a random walk of (*x* + 1)/*b* steps, where *b* ≈ 2.4 ntds is the Kuhn length of single stranded RNA [34, 62]. Since the entropic cost of forming base pairs has already been considered in Δ*S*_stem_, for the purposes of calculating Δ*S*_loops_ we consider stems as rigid rods. This approximation is justified because of the extremely long persistence length of double-stranded RNA (~ 200 ntds [63]) compared to both single-stranded RNA and the length of any stem we consider.

The entropy of a single-stranded region of length *s_i_* is given by *k_B_* log *ω_i_*(*s_i_*), where *ω_i_*(*s_i_*) is the number of ways of arranging the region consistent with the topology of the overall structure. Defining Ω(*s*) as the *total* number of conformations a random walk of length s can take, for a free chain, *ω* = Ω. For structures which include constraints, *ω*(*s_i_*) = Ω(*s_i_*) × *p*(*s_i_*), where *p*(*s_i_*) is the probability that the random walk of length *s_i_* will yield a conformation consistent with the topology of the overall structure being considered. Since free energies are measured relative to the free chain, factors of Ω cancel out in equations for Δ*S*_loops_ (see further discussion in Section S3). The entropy of the single-stranded regions in a given structure is thus given by

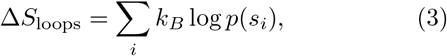

where *s_i_* is the number of nucleotides in the *i*^th^ singlestranded region. The sum is generally over non-independent terms; we will describe how to address these sums via a Feynman diagram-like approach in this section.

As demonstrated in Eq. (3), the physics of the situation are held in *p*(*s*), which is best calculated by considering the end-to-end vector of the random walk undergone by the single-stranded RNA, as

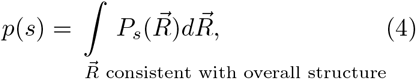

where we define 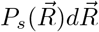 as the probability of a random walk of length *s* to have end-to-end vector 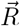:

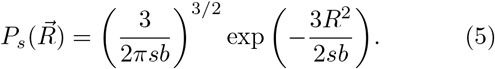

We have assumed *s* ≫ *b* in order to arrive at the Gaussian formula above through the central limit theorem. The mean of the Gaussian is zero by symmetry. In order to find the variance we first consider a single step of length b in three dimensions which has variance in the *x, y*, and *z* coordinates of *b*^2^/3 by symmetry. For a random walk of *N* = *s*/*b* steps, by independence of subsequent steps, the total variance is equal to *Nb*^2^/3 = *sb*/3, leading to Eq. (5).

As described in Section S5 of the supplement, we can systematically consider higher order corrections to Eq. (5) while maintaining its Gaussian nature. Eq. (5) is accurate for non-self-avoiding random walks; self-avoiding random walks cannot be treated analytically in this way. However, for sufficiently short walks, the probability of self-interaction is low. While the accuracy of the assumption *s* ≫ *b* does not always hold in the problems considered, we ultimately find very good agreement between results using Eq. (5) and experiment, and that corrections to Eq. (5) as described in Section S5 are negligible.

In order to demonstrate how Eqs. 3–5 are applied, we first consider the simple hairpin loop. Following Jacobson and Stockmayer [65], we allow that base pairing can occur as long as the two nucleotides are within a small volume 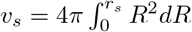 of one another, where *r_s_* roughly corresponds to the bond length.^1^ We assume that *r_s_* is small enough that for all 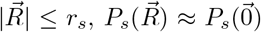. Therefore, Eqs. 3–5 yield

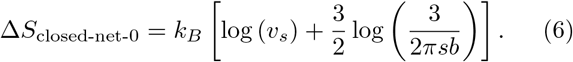

We have called the LHS of the equation *S*_closed-net-0_ (the zero references the lack of stems enclosed by the loop) following [34, 52] (rather than, say, *S*_hairpin_) to emphasize that this formula is applicable to hairpin loops, bulge loops, internal loops, and multiloops – all of which can be thought of as closed loops of RNA. Aside from the appropriate inclusion of *v_s_* terms to account for the finite and variable width of RNA stems, RNA stems are treated as having negligible width by performing the approximation 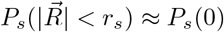.

We estimate *v_s_* by fitting experimental measurements of the entropy of hairpin loops of variable lengths to Eq. (6). Although Eq. (6) implies that the entropy of a hairpin should increase monotonically as a function of its length, the experimental measurements are nonmonotonic, and their nonmonotonicity exceeds the error bars [64]. This non-monotonicity may be due to enthalpic effects [66] which were neglected in our analysis following Ref. [25]. Nevertheless, Fig. 2 shows that Eq. (6) gives a reasonable fit to the experimental data with *v_s_* = 0.0201 ± 0.0036 ntds^3^.^2^ If one ignores all angular dependences of bond formation, this leads to a naive underestimate of the length of a hydrogen bond of 0.56 Å, which nonetheless is well within an order of magnitude of the true length of hydrogen bonds.

**FIG. 2:**
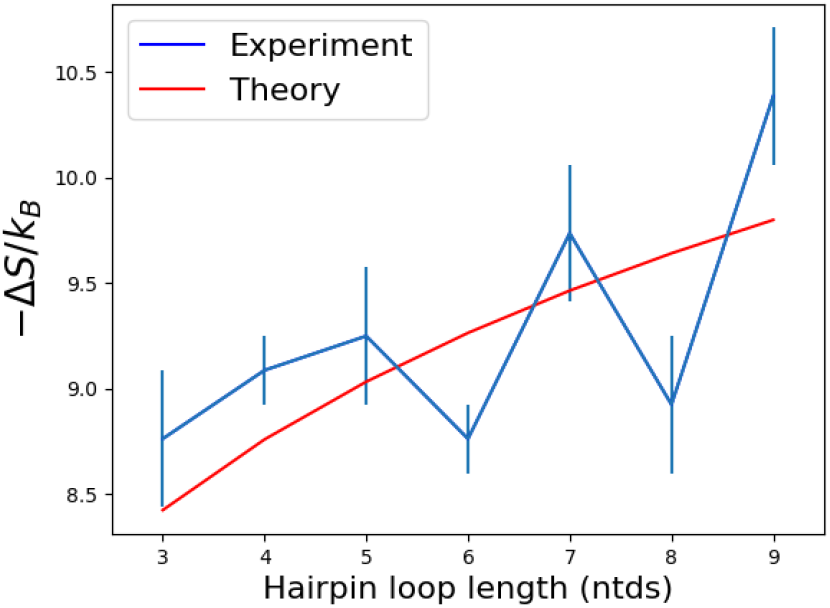
*v_s_* estimated from experimental data. Experimental estimates for the free energy of hairpin loops of length s from Table 1 of Ref. [64] were converted to entropy estimates (blue points and error bars) by assuming Δ*H* = 0 as in Ref. [25]. These data were fit to Eq. (6), yielding an estimate of *v_s_* = 0.0201 ± 0.0036 ntds^3^.

Finally, we consider pseudoknots. To calculate the entropy of a pseudoknot of arbitrary complexity we invent a novel graph formulation inspired by Feynman diagrams from quantum field theory. First, the RNA structure being considered is translated into a graph. Nodes are used to represent the two end points of a stem, and two types of edges represent single- and double-stranded RNA.

Defined in this way, the graph of the RNA structure directly represents the integrals necessary to compute its entropy. The positions of the nodes are integrated over all of space, while the constraints of the structure are included in the integrand: a double-stranded edge of length *l* between nodes *i* and *j* leads to a term 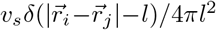, and a single-stranded edge of length *s* between these nodes leads to a term 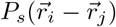 in the integrand. Note that two bonded nucleotides in isolation are considered a stem of length *l* → 0.

As a concrete example, we consider the canonical H-type pseudoknot, an instance of which is shown in Fig. 3A (LHS). As we described, its conformational entropy can be calculated by translating the structure into a graph (Fig. 3A RHS), where each node represents the edge of a stem; blue edges represent regions of doublestranded RNA of length *l_i_*; red edges represent regions of single-stranded RNA of length *s*_i_. For example, here, *s*_3_ = 5 ntds, and *l*_1_ =3 ntds. We set the origin of our coordinate system to node 0 and call the distance between node *i* and the origin *r_i_*. Integrating over the possible placements of nodes 1-3 (while including the constraints of the structure in the integrand as described previously) we obtain the following Gaussian integral formulation of the entropy:

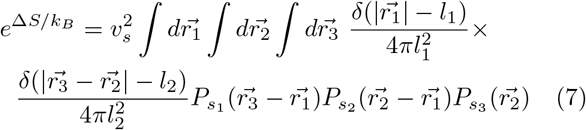

where using the assumption *s* ≫ *b*, we allow the integrals to extend over all of space. A more comprehensive derivation of this formula, including the origin of the *v_s_* terms, can be found in Section S4. This integral can be calculated analytically (Sec. S5) [34].

**FIG. 3:**
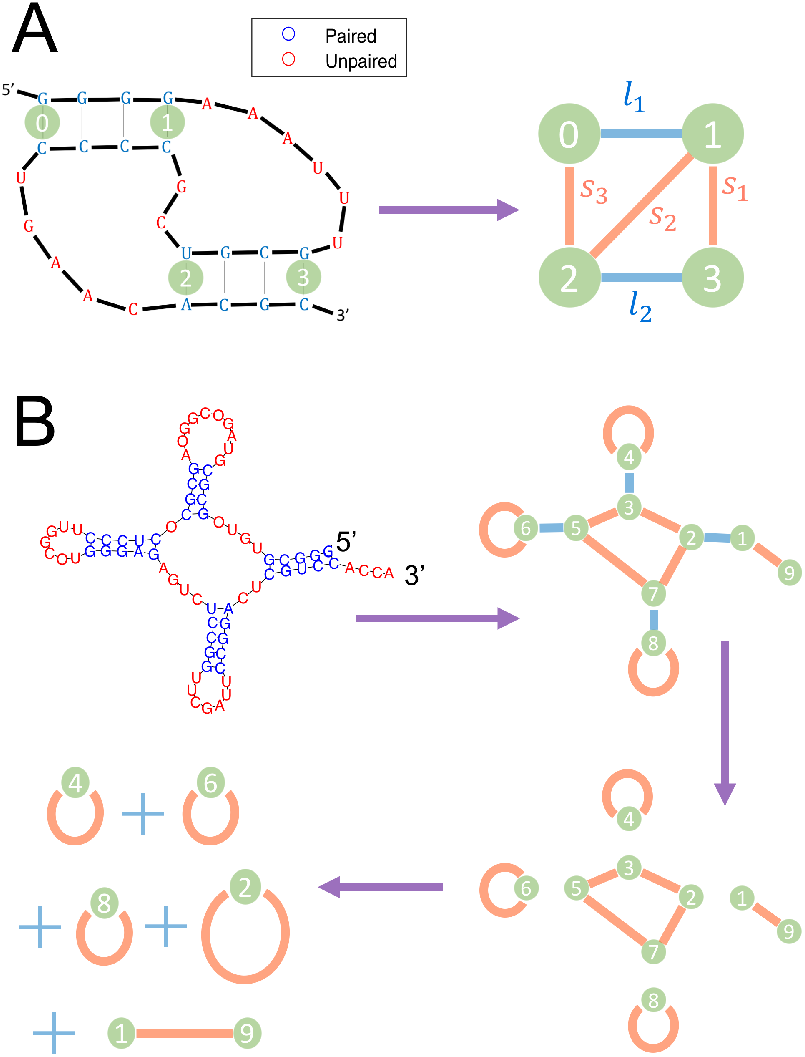
RNA Feynman Diagrams. (**A**): **The Canonical Pseudoknot** An instance of the canonical H-type pseudoknot. Bold lines represent the RNA backbone; thin lines represent Hydrogen bonds. The entropy of this structure can be calculated by converting it to a graph format as shown in RHS of panel. The nodes of the graph represent the first and last base pairs of each stem, and two types of edges represent single- and double-stranded RNA. The graph directly represents the integral in Eq. (7). (**B**): **Graph Decomposition**. The entropy of a sample RNA structure (top left) can be computed by converting the structure to a graph as defined in the text (top right). The graph directly represents the integrals necessary to compute the entropy. Separable integrals are represented by graphs which can be disconnected by the removal of any one edge (bottom right). Thus, once appropriate factors of Vs are included (one for each stem in the original structure), the entropy of the structure in question is given by (bottom left) the sum of four closed-nets-0 (originating from the three hairpins and multiloop) and one open-net-0.

Graphs that can be disconnected by the removal of any one edge correspond to separable integrals, and thus to distinct motifs in the RNA structure. The decomposition of a structure into its component graphs is depicted in Fig. 3B for a classical cloverleaf RNA. The RNA in question decomposes into four instances of closed-net-0 (originating from the three hairpins and multiloop) and one instance of an open-net-0, or free chain (which by definition does not affect the entropy). As shown in the figure, once appropriate factors of *v_s_* are included in the integrals (one for each stem) the stems can be treated as having negligible width; thus, nodes which can be removed without changing the topology can be removed in the graph decomposition process. See Section S4 for further discussion.

In Fig. S2 we display all possible graphs of up to two stems and their respective RNA structures. As in Fig. 3, single-stranded edges are displayed with red; doublestranded with blue. For each graph, the integral formulation of its entropy is displayed in the figure alongside what it evaluates to.

## IV. COMPARISON WITH PUBLISHED TOOLS

We use experimentally determined structures to compare the predictions of our model with other current methods; results are shown in Fig. 4. For sequences of length ≤ 80 ntds from the RNAStrand [53], PseudoBase++ [54], and CompraRNA [55] databases (186 non-pseudoknotted structures with 58 different topologies; 235 pseudoknotted structures with 52 different topologies) which had a sequence dissimilarity ≥ 0.2 (using Jukes-Cantor) we measured the number of base pairs correctly predicted by our algorithm’s MFE structure compared to fourteen other current algorithms. Seven of these cannot predict pseudoknots and serve as useful benchmarks for the non-pseudoknotted results, (detailed methods in Section S1).

While the entropy model presented here can give an integral expression for arbitrarily complex pseudoknots, the integral may need to be solved numerically for sufficiently complex structures. For this large-scale comparison we disallowed pseudoknots more complex than those displayed in Fig. S2, and our algorithm therefore did not require any numerical integration. We similarly disallowed parallel stems which can be stable in neutral and acidic pH conditions [73]. We also set the minimum stem length for each sequence (*m*) to the minimum value it could take such that the total number of possible stems is less than 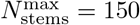. These choices were all made to speed up computation time; each sequence took between several seconds and ~ an hour to run on a MacBook Pro 2012 laptop. Details of the computation time of our algorithm can be found in Fig. S4.

**FIG. 4:**
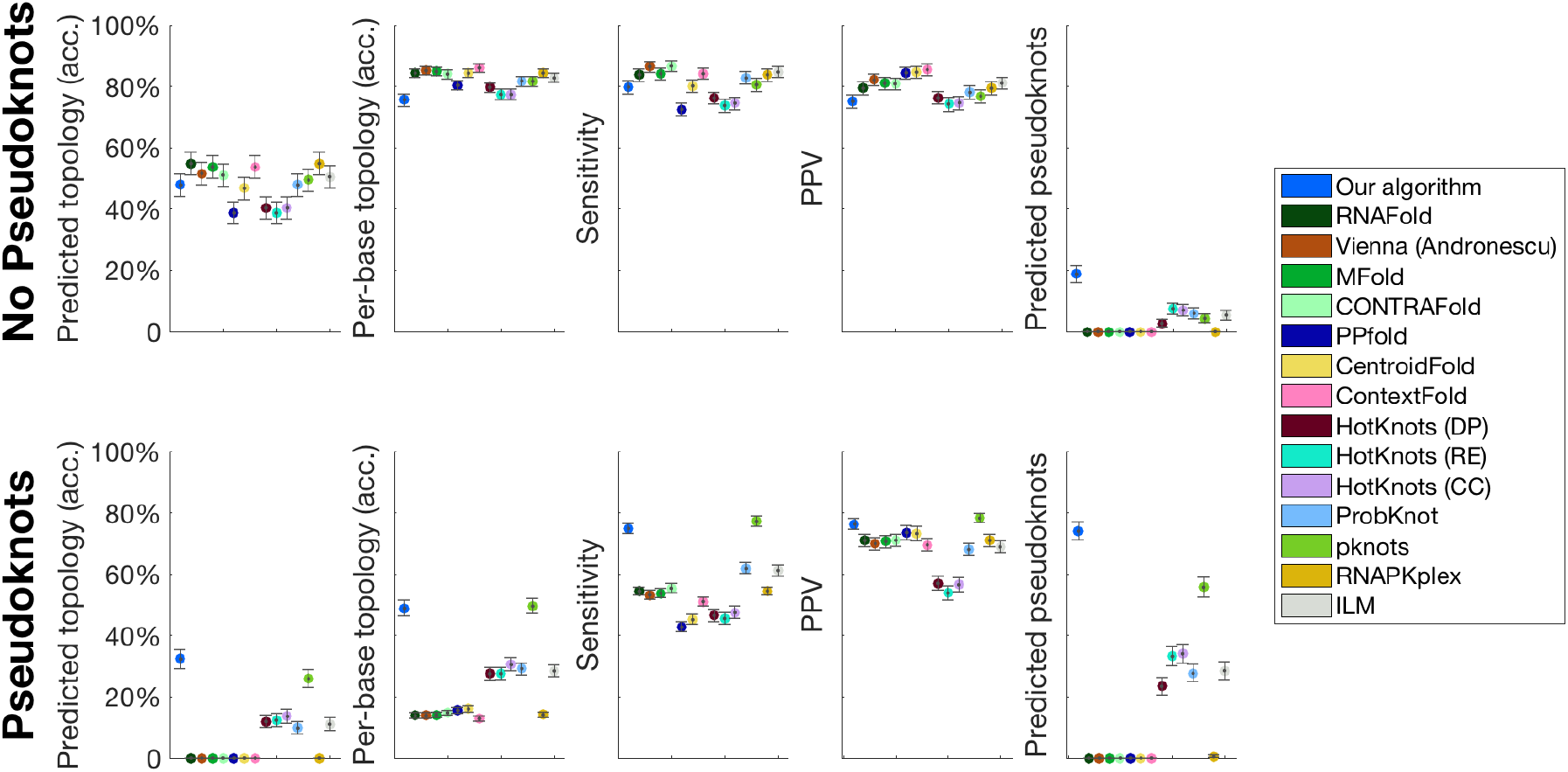
Summary statistics for comparison to other prediction tools. To assess the relative success of our algorithm, we compare its performance to that of 14 other current prediction tools: RNAFold [29, 67], ViennaRNA (Andronescu parameters) [68], Mfold [28], CONTRAFold [69], PPfold [70], CentroidFold [71], ContextFold [72], HotKnots (Dirks-Pierce parameters), HotKnots (Rivas-Eddy parameters), HotKnots (Cao-Chen parameters) [51], ProbKnot [37], pknots [39], RNAPKplex [29, 67], and ILM [35]. We measure sensitivity, PPV, the fraction of topologies predicted correctly by the MFE structure, the average per-base topology accuracy (defined in the main text), and the proportion of the time the MFE structure contains a pseudoknot. We separate the results for sequences which form into pseudoknots and those which don’t. Error bars show the standard error. Despite the fact that our algorithm requires only two parameters to describe the entropy of any arbitrary secondary structure (at least an order of magnitude – and often several – fewer than the other algorithms tested against), and that the parameters were trained on non-pseudoknotted structures, our algorithm outperforms the other algorithms tested in predicting pseudoknotted structures, and performs on par with them in predicting non-pseudoknotted structures. See main text for further discussion.

While these practical constraints were chosen to speed up the computation time, they also led to errors in the algorithm’s predictions. 64 of the tested pseudoknots were topologically more complex than any of those presented in Fig. S2. Furthermore, 33 of the non-pseudoknotted sequences tested (and 8 of the pseudoknotted) include base pairs outside of those allowed by the algorithm (A·U, G·C, and G·U). Removing such structures from our comparison analysis leads to our algorithm performing even better compared to current tools (see Fig. S3).

Further errors were due to our choice of *m*, which was not optimized and was too high compared to the length of the shortest stem in the experimental structure for 58 non-pseudoknotted cases and 54 pseudoknotted. By changing 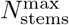 from 150 to 200, these numbers decreased to 46 for both pseudoknotted and non-pseudoknotted sequences, but the results for 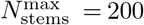 were practically identical to the results of Fig. 4 (see full results in Supplementary Table 1). For 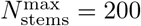, the computation time was increased significantly (to ~ 17 hours for one sequence).

The sensitivity (*TP*/*TP* + *FN*) and PPV (*TP*/*TP* + *FP*) of our algorithm were measured to be 0.80 and 0.75 for the non-pseudoknotted cases, and 0.75 and 0.76 for the pseudoknotted cases, respectively. Our algorithm outperformed all other prediction tools tested for the prediction of pseudoknots, and on par with other tools in the prediction of non-pseudoknotted sequences. The full results can be found in Supplementary Table S1.

While sensitivity and PPV are the most common metrics used to establish the success of an RNA prediction algorithm [74], we sought to develop a test that measures success on the scale of the full RNA, rather than on the scale of individual base pairs. To this end, we measured how frequently each algorithm was able to correctly predict the topology of the experimentally measured structure, where the topology of a structure is defined by its graph (Section III). We found for our algorithm that the experimental topology is within the top 1, 5, and 10 topologies at frequencies of (49%, 65%, and 70%) for non-pseudoknotted structures, and (34%, 59%, and 62%) for pseudoknotted, demonstrating a sharp increase between top 1 and top 5, and a plateau between top 5 and top 10.

Considering whether an algorithm correctly predicts the full topology can lead to errors arising from small variations in structure. For example, the opening of a single bond on the edge of a stem can lead to a different topology as we’ve defined it, if that stem includes one of the ends of the molecule. In order to arrive at a per-base measure of topology, we consider for each bond along the RNA backbone to which of the minimal graphs of Fig. S2 it belongs. For example, the bond between the second and third nucleotides of Fig. 3A belong to a stem of an open-net-2a graph. We then measure for each sequence the fraction of correct per-base topology predictions made by each algorithm’s predicted MFE structure. We find that our algorithm averages an 76% per-base topology prediction accuracy for non-pseudoknotted sequences, and a 49% accuracy for pseudoknotted.

Finally, we compare how frequently each algorithm predicts an MFE structure containing a pseudoknot. Our algorithm correctly predicted 174/235 pseudoknots among the pseudoknotted cases, far more than any other algorithm tested. However, it also erroneously predicted 35/186 incorrect pseudoknots among the non-pseudoknotted cases. We have found that the probability of predicting pseudoknots can be significantly decreased with minor changes in the Turner parameters energy function, and these parameters may need to be re-examined in order to be used most effectively with the entropy model presented here.

Our algorithm also provides the probability of folding into a pseudoknotted structure for each sequence. These data for the 421 sequences tested are presented in Fig. 5. Each datapoint represents a different sequence and the total probability calculated of that sequence folding into a pseudoknotted structure. For figure clarity, a lower bound of pseudoknot probability was set at 2 × 10^−10^.

**FIG. 5:**
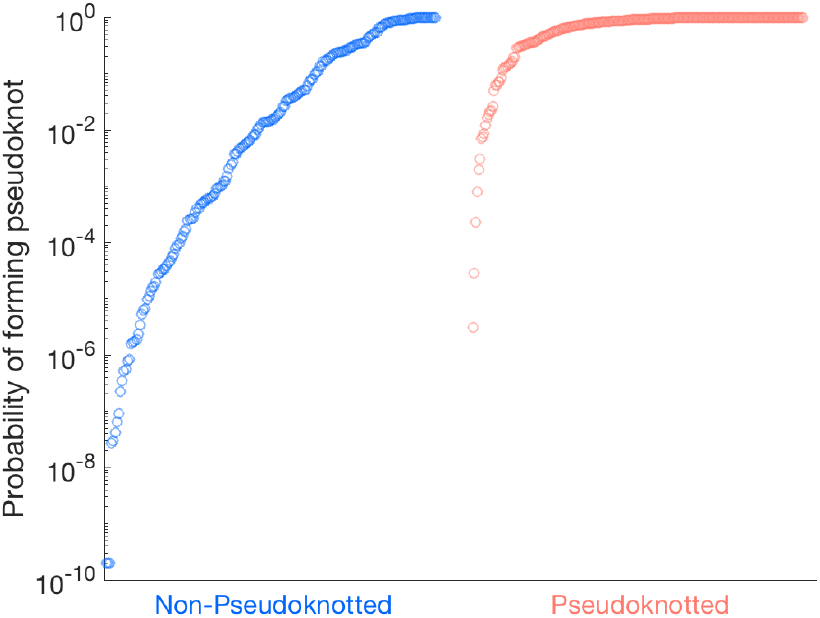
Probability of folding into a pseudoknot. The predicted probability of each of the 421 sequences tested folding into a pseudoknot is presented. Of these sequences, 186 were experimentally found not to form pseudoknots (blue) and 235 were found to form pseudoknots (red). Our algorithm successfully predicts pseudoknots forming in the latter category far more frequently than in the former. For figure clarity, a lower bound of pseudoknot probability was set at 2 × 10^−10^.

The algorithm’s predictions for the six longest RNA molecules less than 89 ntds in length from the Pseudobase++ database are presented in Fig. 6. We considered only those sequences whose structure was directly supported by experiments and which could be decomposed into the minimal topologies shown in Fig. S2. We display the experimental structure (green background) alongside the MFE predicted structure (light blue background) and the top six predicted topologies (out of several hundred, depending on the sequence; dark blue) where the experimental topology is highlighted (purple). RNA secondary structure was plotted using the PseudoViewer package [83]. Our results demonstrate successful predictions even for long pseudoknotted sequences, especially in terms of the predicted topology. Detailed methods are provided in Section S1.

**FIG. 6:**
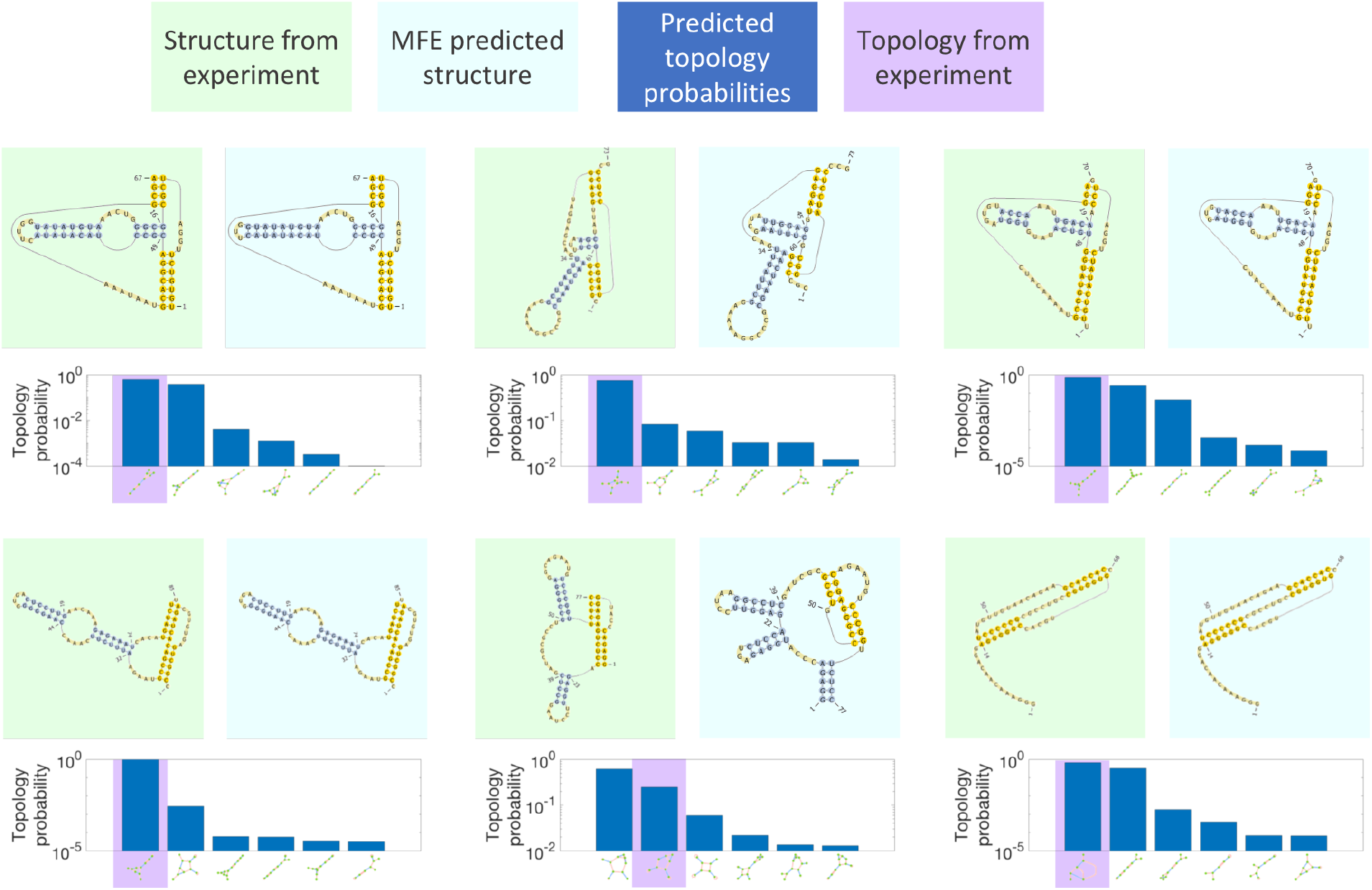
Comparison to experiments for long sequences. Six long sequences were chosen from the Pseudobase++ database as described in the main text. The sequences are derived from (starting from the top left and moving across): tobacco mosaic virus [75–77], *Bacillus subtilis*, [78], tobacco mild green mosaic virus [76, 79], *Bacilis subtilis* [80], Giardiavirus [81], and Visna-Maedi virus [82]. We show the experimental structure (green background) and the MFE predicted structure (light blue background) plotted using the PseudoViewer software [83]. We also display the top six topologies (out of several hundred, depending on the particular sequence) and their respective predicted probabilities, with the topology corresponding to the experimental structure highlighted in purple. Overall, our results demonstrate successful predictions even for these long pseudoknotted sequences, especially in terms of the predicted topology.

## V. DISCUSSION AND CONCLUSIONS

The accurate prediction of the ensemble of secondary structures explored by an RNA or DNA molecule has played a major role in shaping modern molecular biology and DNA nanotechnology over the past several decades. In this work, we showed that the modern ubiquity of extremely powerful computers can be used alongside novel polymer physics techniques to completely enumerate the free energy landscape of an RNA molecule including complex pseudoknots. This NP-complete algorithm can be used to tackle even relatively long (~ 90 ntds) RNA sequences, and aside from the enumeration procedure (which is relatively fast for long sequences; see Fig. S4) is easily parallelizable.

Remarkably, the entropy model discussed in this work requires only two parameters – orders of magnitude fewer than other current algorithms – corresponding to clearly measurable physical quantities. Despite this, and despite the fact that all parameters used in our model were derived using experiments on non-pseudoknotted RNA, our algorithm is more successful in predicting pseudoknotted structures than any of the other algorithms tested, and on par with all predictors tested in predicting non-pseudoknotted structures. Although we have not done so in this work, we expect that our results can be even further improved by optimizing the energy function given the entropy model presented here. The success of our algorithm is particularly notable given that the entropy model developed in this work can be used to address any RNA secondary structure regardless of complexity.

The algorithm presented here can also be easily generalized to probe multiple interacting strands (see discussion in supplement). The sequences considered can be any combination of DNA and RNA; their identities affects the energy parameters of the model which have been previously tabulated, and to a lesser extent the two entropy parameters (*b* and *v_s_*).

Our finding that the integral formulation of the entropy of arbitrary complex RNA secondary structures can be represented graphically is reminiscent of Feynman diagrams in quantum field theory. The topologies defined by these graphs can also serve as useful biological constructs to group similar RNA structures together. The depiction of RNA structure as a graph has played an important role in the prediction of RNA secondary structure [84–87], as well as in the search for novel RNAs [88, 89], and the description of similarity between RNA structures [90–93] which is especially useful in the study of the effects of mutations [94, 95]. A common approach among these graphical depictions of RNA has been to represent loops (e.g. hairpins, internal loops, etc.) as verticies and stems as edges [88, 92, 93]. However, this depiction of RNA does not always distinguish between pseudoknotted and non-pseudoknotted structures [88]. Other approaches have represented each nucleotide as a separate node and bonds (either hydrogen or covalent) as edges [89, 91]; while useful in many contexts (for example, secondary structure visualization), this approach does not have the benefit of coarse-graining to group similar structures as the same graph [90]. Our approach, described in Section III, can be viewed as a middle ground and may be useful in the contexts described previously.

## VI. ACKNOWLEDGEMENTS

We thank Elena Rivas and Yohai Bar Sinai for fruitful discussions. This research was funded by the National Science Foundation through the Harvard Materials Research Science and Engineering Center Grant DMR-1420570, DMREF Grant DMR-123869 and ONR Grant N00014-17-1-3029. OK acknowledges support from an NDSEG fellowship and Molecular Biophysics Training Grant NIH/NIGMS T32 GM008313 (PI: James M. Hogle). M.P.B. is an investigator of the Simons Foundation.

## Supporting Information

The supporting information is divided into several sections. In Section S1 we detail the methods used to compare our algorithm’s performance to other current models. In Section S2 we discuss how our algorithm can be easily generalized to probe multiple interacting strands including any combination of DNA and RNA. In Section S3 we give a further discussion of the Ω(*s*) term defined in Section II. In Section S4 and Fig. S1 we provide a more complete derivation of Eq. (7). In Section S5, we show how to analytically calculate the integrals in Eq. (7). In Section S6 we derive the higher-order corrections to Eq. (5).

In Fig. S2 we display all possible graphs of up to two stems and their respective RNA structures along with the integral formulation of their entropies and their evaluated forms. In Fig. S3 we discuss how our algorithm compares to state-of-the-art prediction tools (the analogue of Fig. 4) when restricting ourselves to structures allowed by the chosen constraints on our algorithm. Finally, in Fig. S4 we show how our algorithm’s properties scale with the length of the sequence for random sequences between 10 and 21 ntds in length.

### S1. DETAILED METHODS FOR COMPARISON WITH OTHER PREDICTION TOOLS

In order to compare the sensitivity and PPV of different prediction tools, we considered the base pairs present in the experimental structure and in each algorithm’s MFE structure. Base pairs present in both were labeled as true positives (*TP*), base pairs present in the predicted algorithm were labeled as false positives (*F P*) and those present in the experimental structure but not the predicted MFE structure were labeled as false negatives (*FN*). In order to compare different metrics we use the summary statistics of sensitivity (*TP*/*TP* + *FN*) and PPV (*TP*/*TP* + *FP*). PPV is a more useful metric for RNA structure prediction algorithms than specificity because the definition of true negatives is unclear when considering base pairs.

The sequences tested were downloaded from the Pseudobase++, RNAStrand, and CompraRNA PDB databases. We constrained database searches to return results only for sequences of length ≤ 80 ntds. We further restricted the search of the RNAStrand database to only include sequences where all nucleotides were known, and to not include fragments, multiple strands, or duplicates. We removed all sequences that had hairpins of under 3 ntds. Finally, we compared the sequence similarity of the sequences derived and kept only sequences with ≥ 0.2 Jukes-Cantor sequence dissimilary measured using the MatLab command `seqpdist`. The Jukes-Cantor distance between two sequences is defined as

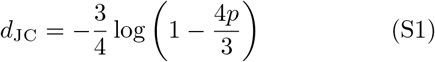

where *p* is the fraction of sites which differ between the sequences after they have been aligned. By imposing *d*_JC_ ≥ 0.2 we impose a constraint that *p* > 0.17.

We assumed *T* = 300*K* for all predictions.

In order to speed up computation for longer sequences, we set the parameter *m* describing the minimum number of consecutive base pairs in a stem to the minimum value it can take such that the total number of possible stems is less than 150. This latter parameter was chosen arbitrarily and is likely not optimized; however, changing it to 200 had no significant effect (see data in Supplementary Table 1). This resulted in *m* = 1 for 22% sequences, *m* = 2 for 33% of sequences, *m* = 3 for 23%, *m* = 4 for 20%, and *m* = 5 for nine sequences. Changing the maximum total number of possible stems to 200 resulted in *m* = 1 for 34% sequences, *m* = 2 for 29% of sequences, *m* = 3 for 22%, *m* = 4 for 15%, and *m* = 5 for one sequence.

Our algorithm can enumerate and calculate the entropies of both parallel and antiparallel stems. (An antiparallel stem is a list of consecutive base pairs of the form [*i* · *j*, (*i* + 1) · (*j* − 1), (*i* + 2) · (*j* − 2)…], while a parallel stem has the form [*i*·*j*, (*i* + 1) · (*j* + 1), (*i*+2)·(*j*+2)…].) Parallel stems are disallowed in non-pseudoknotted structures, and are stabilized at certain pH levels. We disallowed parallel stems in our calculations.

As part of the enumeration procedure, we created a compatibility matrix *C_p,q_* detailing the compatibility of structures *p* and *q* (structures *p* and *q* are compatible if they do not share any nucleotides). In practice, since there are some structures whose entropies we have not analytically derived, we found it useful to also construct three- and four-dimensional matrices *C*_3_ and *C*_4_ which define three- and four-way compatibility, in order to exclude most such structures at this stage.

In order to compare topologies, we measure whether the eigenvalue spectra of the two matrices defining the bonds between each node are equal (two matrices are needed because there are two types of bonds). This method is guaranteed to correctly identify graph isomorphisms in all cases but may have false positives. We have found no evidence of false positives in all cases tested (compared against the MatLab `isisomorphic` command).

For the analysis in Fig. 6 we also set *m* > 1 to speed up computation. Starting from the top left and going across, we set *m* = (4, 3, 3,4, 4, 4). We also disallowed parallel stems in order to speed up the computation.

### S2. PROBING MULTIPLE INTERACTING STRANDS

The algorithm presented here can also be easily generalized to probe multiple interacting strands, using only one further parameter which has been previously studied to define the free energy cost of forming a duplex [96, 97]. Following Ref. [28] we concatenate the two (or more) sequences, separated by a number of inert nucleotides which serve as a placeholder and which are removed before free energy calculations are implemented.

The algorithm described here can be equally well-applied to DNA strands by using the parameter sets from the SantaLucia laboratory [98–102]. In addition, our algorithm can probe DNA-RNA bonds using the parameter sets from Refs. [103, 104], and interpolating between the DNA and RNA cases for those parameters that have not yet been tabulated from experimental data. The inclusion of DNA strands may require slight modification to the two entropy parameters (*b* and *v_s_*) which are based on data from RNA experiments.

### S3. DISCUSSION OF Ω(*s*)

In Section II, we defined a parameter Ω(*s*) to be the total number of conformations a random walk of length *s* can take. Ω(*s*) has the property that Ω(*s*_1_)Ω(*s*_2_) = Ω(*s*_1_ + *s*_2_).

For a free chain of length *s*, 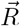 can take on any value, as long as 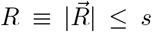. Taking the limit *s* ≫ *b* (so that the integral extends over all of space), and using the normalization of 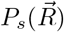, we find *S*_free_ = *k_B_* log Ω(*s*), as expected from the definition of Ω(*s*). Therefore, in order to calculate changes in free energy compared to *S*_free_, we simply omit Ω(*s*) terms from our formulae.

To be more precise, this argument directly demonstrates only why Ω(*s*) terms should cancel out for nonbase-paired RNA. However, it motivates us to consider the experimentally measured entropy of each base pair as being multiplied by a factor of Ω(2) for the two nucleotides comprising it. Including such terms, all factors of Ω drop out of expressions representing physical results. We therefore can compute expressions for Δ*S*, the difference in entropy between a given structure and a free chain, by omitting factors of Ω from the relevant formulae.

### S4. DERIVING EQ. 7

In this section we more fully detail the steps leading to Eq. (7), the entropy of the RNA structure depicted in Fig. S1A.

We start by treating each nucleotide as its own node, subject to the constraint that the distance between nucleotides is given by *a* = 0.33 nm. Writing such an expression is cumbersome, but because of the property of 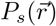 that 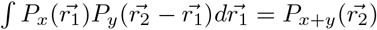, we can simply integrate over all nodes not at the edges of stems.

The full expression for the entropy of this graph is thus given by

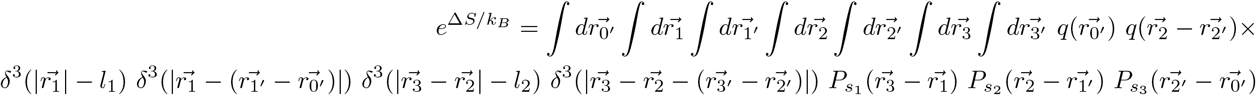

which is depicted graphically in Fig. S1B. We are using *δ*^3^(|*x*| – *a*) to signify

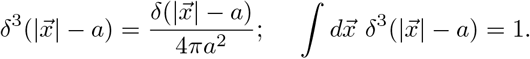

*δ*^3^(|*x*| – *a*), like 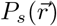, has units of inverse volume.

Vectors are defined relative to the origin where node 0 is placed (i.e. 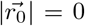). There is no integration over r0 because such an integral would cancel out with the corresponding term in *S*_free_, and thus disappear in the formula for Δ*S*.

One can check that introducing a new node representing any nucleotide in the structure (say a node on the edge between nodes 0 and 3) does not affect the result.

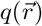 is defined as the probability of a nucleotide located a vector 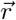 from the origin to be bonded to a nucleotide located at the origin (assuming the two nucleotides are complementary). If following Ref. [65] we wish to include an upper bound for the bond length, *r_s_*, 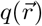 becomes a Heaviside Θ function. Integration over *q* leads to the definition of 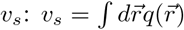.

Only two factors of *q* are present, as opposed to one factor for each base pair in the structure, because we take the entropy of stems into account separately. For this expression, we treat stems as rigid rods; while the rods have variable and finite width (corresponding to the property that nucleotides do not need to be at a precise separation in order to bond), they cannot be thicker on one end than the other, since including such possibilities would overcount the entropy of the stem. Our expression thereby has the property that it is invariant if we also integrate over two nodes representing two arbitrary base pairs (say, one on the stem between node 0 and node 1, and one between nodes 0′ and 1′). The choice of which bonded nodes on each stem to put in the argument of *q* is arbitrary, but there is only one bonded node (and therefore one *q* term) for each stem.

We make progress by assuming that because of the *q* terms and delta functions, nodes representing nucleotides which are bonded are located close enough that the vector 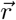 between them can be approximated as having zero length within the context of the terms 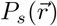.

We therefore approximate our formula as

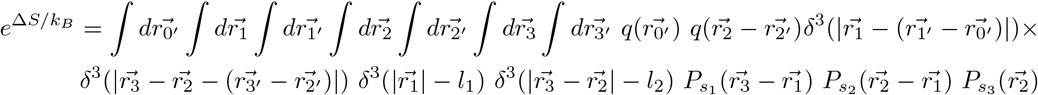

By employing transformations as in Section S5 (e.g. 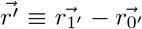), the four integrals over the primed nodes become two integrals over delta functions (which give unity) and two over the *q* terms. The latter two become two factors of *v_s_*, and we arrive at Eq. (7).

**FIG. S1:**
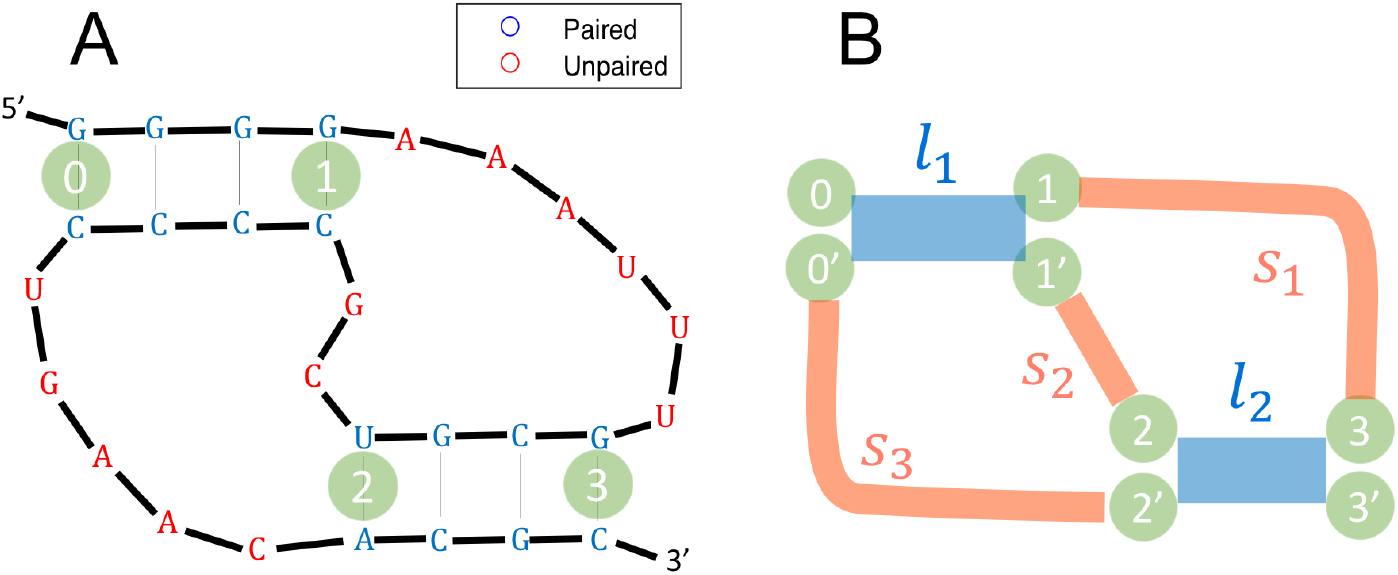
A preliminary description of an H-type pseudoknot. **A**: An instance of the canonical H-type pseudoknot, reprinted from Fig. 3. **B**: A preliminary version of the graph representing its entropy. In Sec. S4 we demonstrate that this graph is equivalent to that shown in Fig. 3A.

### S5. PERFORMING THE GAUSSIAN INTEGRALS

The method of performing the Gaussian integrals of Eq. (7) can be generally applied to the calculation of the entropies of other pseudoknots, and so we describe it in detail here.

Eq. (7) is given by

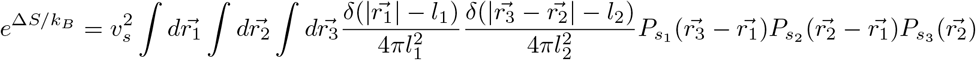

We start by utilizing our approximation that the integrals extend over all of space to rewrite 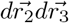 as 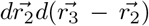, and we rewrite all instances of 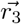 as 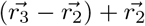.

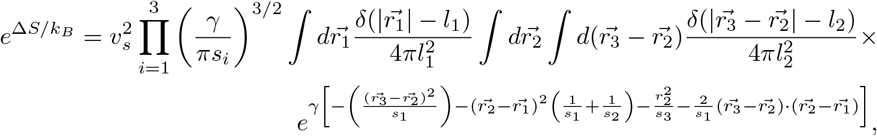

where for notational convenience have defined a parameter *γ* = 3/2*b*.

To do the 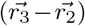 integral, we convert to polar coordinates such that 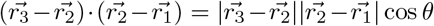. Performing the integral yields

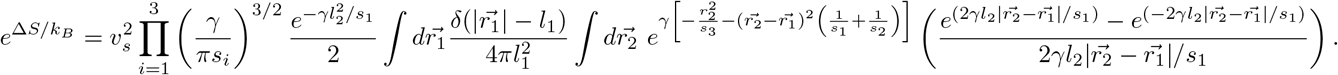

We now use the same trick from before to rewrite 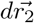 as 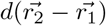, and rewrite each instance of 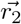 as 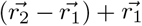. As before, 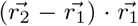 becomes 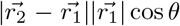. Denoting (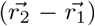 as 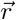 and doing the integral over *r*_1_ after performing this transformation yields

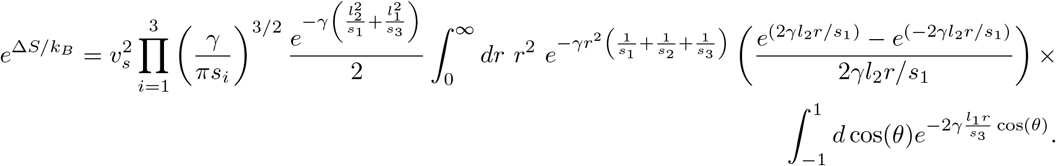

Finally, we perform the integrals remaining to arrive at

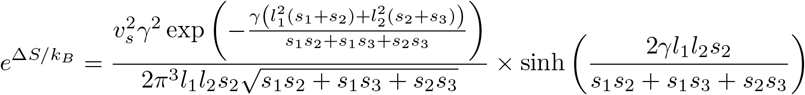

where sinh is the hyperbolic sine function. This formula is equivalent to the one presented without proof in Ref. [34].

### S6. HIGHER ORDER CORRECTIONS TO ENTROPY

Eq. (5), which gives the probability of a random walk of length *s* to have end-to-end distance 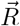, is valid only in the limit of *R* ≫ *b* (where we’ve denoted 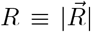). For shorter walks, the Central Limit Theorem no longer holds. In this section, we show a systematic approach to deriving higher-order corrections to the probability distribution given by Eq. (5). The approach taken here is based on a textbook by Ariel Amir (to be published).

We consider *n* steps in three dimensions, where each step is taken to be of length *b* with equal probabilities in all directions. Thus, *s* = *nb*. The probability distribution for where a walker will be after *n* = 1 steps is given by 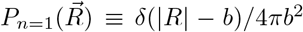. After two steps, the probability distribution for where the walker will be is given by

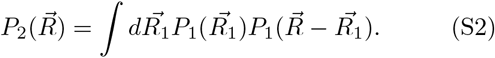

The form of Eq. (S2) is that of a convolution of 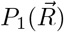 with itself. In order to iterate many convolutions easily, we move to Fourier space, since the Fourier transform of a convolution is the product of Fourier transforms. Fourier transforming 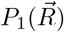 yields its characteristic function: 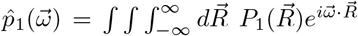, which simplifies to

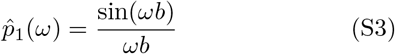

which only depends on 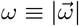.

In order to iterate *n* convolutions in real space, we can simply take the *n*^th^ power of the Fourier transform, finding

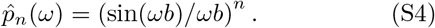

Taking the inverse Fourier transform, we find

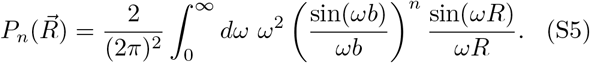

At this point, we use our assumption that *n* is large. This formula tends to zero for large values of *ωb*, and we therefore Taylor expand the sin function for small *ωb*. If we take only the first two terms of this series, we would arrive at Eq. (5); we therefore take the first three terms to get the first correction to Eq. (5). Higher-order corrections can be found by simply taking more terms of the series. Eq. (S5) thus becomes

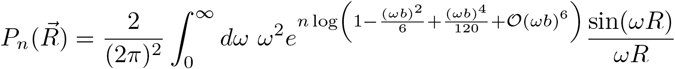

Next, we Taylor expand the logarithm and write the sin as a sum of exponentials. Since the two terms in the sum are identical under the exchange *ω* → −*ω*, we combine them into one term by changing the lower limit of integration to −∞.

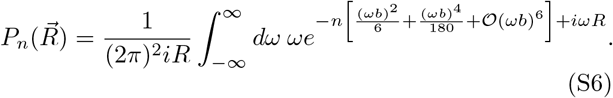

If we didn’t have the quartic term, this integral would be Gaussian and would result in Eq. (5). However, if we keep this term, the integral is no longer solvable analytically. We proceed by setting

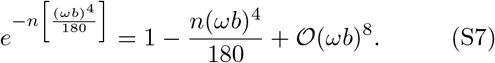

As is apparent, the finite truncation of this series results in corrections of higher order than the truncation of the series for sin(*ωb*) or of the logarithm above.

Using this series expansion, Eq. (S6) becomes a Gaussian integral, which can be solved analytically to yield

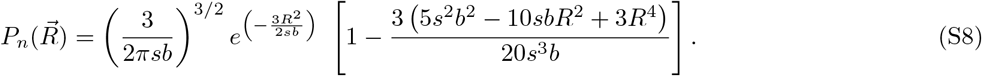

where we’ve replaced *n* by *s*/*b*.

One of the essential properties of 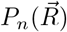 for our formalism to function is that 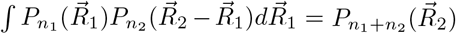. One can check directly that this holds for Eq. (S8). Keeping only first-order correction terms, and defining 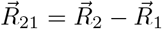,

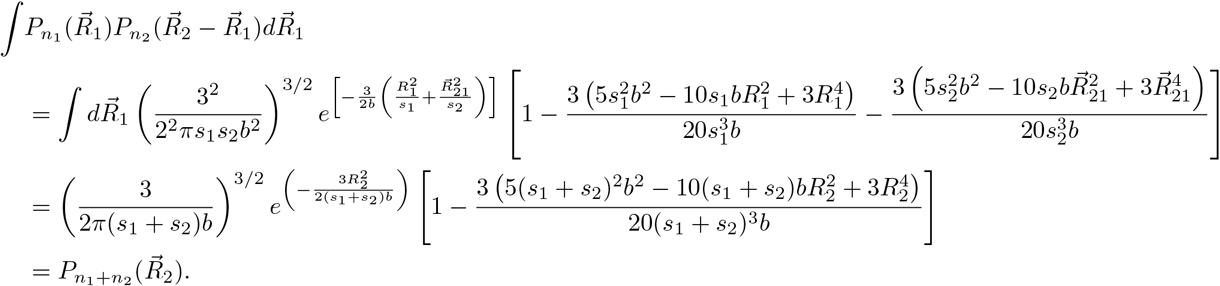

**FIG. S2:**
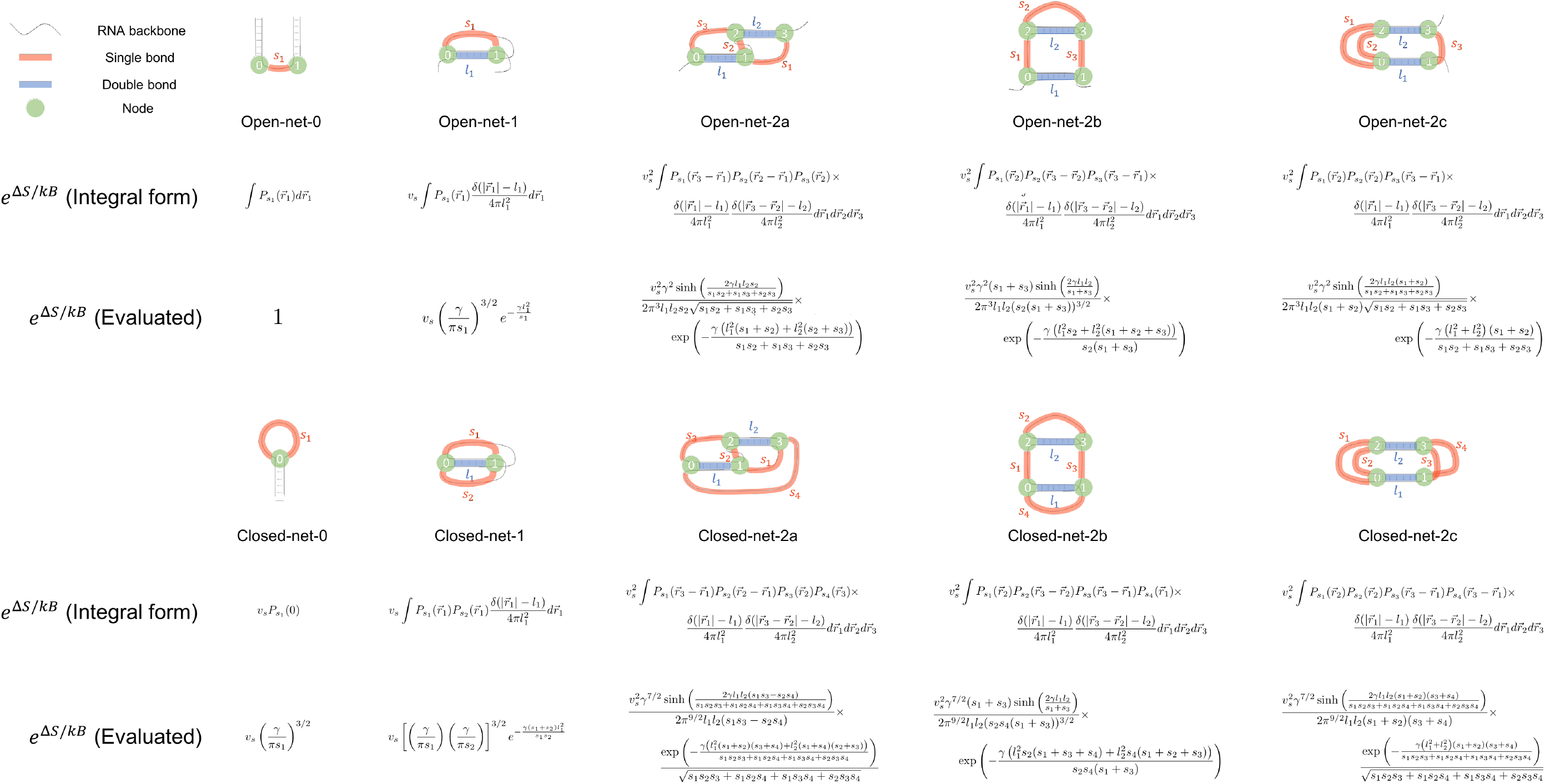
Graphs of simple RNA structures. The 10 graphs with at most two regions of double-stranded RNA and their corresponding RNA backbones are displayed alongside integral and evaluated expressions for the entropy of each graph. Note that stems shown as parallel could be antiparallel if the system considered is comprised of more than one strand. See Fig. 2 of Ref. [52] for comparison.

**FIG. S3:**
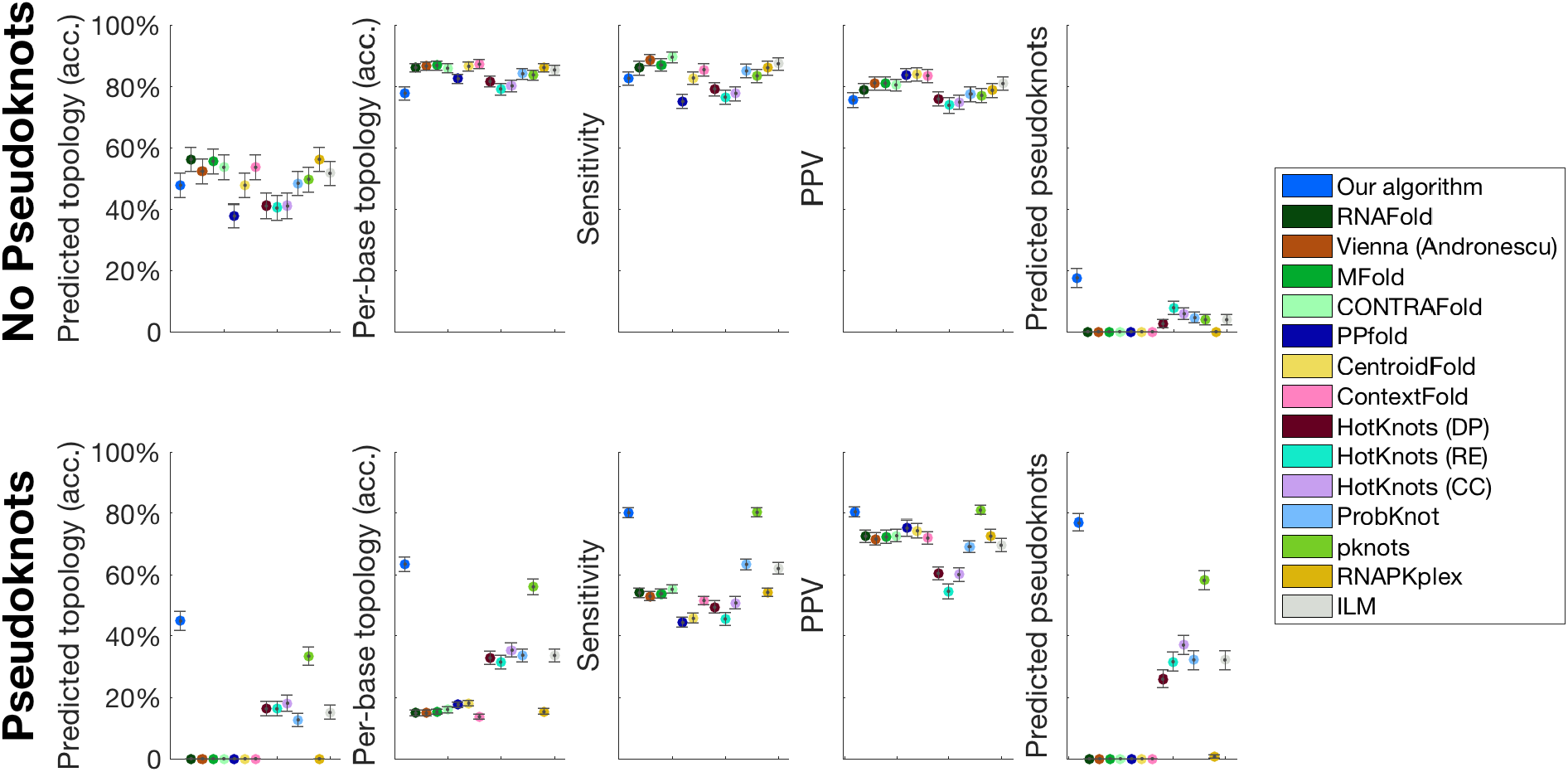
Results only including sequences whose structure our algorithm could have predicted. We consider only the 153 non-pseudoknotted and 165 pseudoknotted sequences whose structures do not include base pairs or topologies disallowed by our algorithm. In this case, we predict the correct topology with 49% (47%) accuracy for non-pseudoknotted (pseudoknotted) structures. This number increases to 62% (82%) and 67% (85%) for top-5 and top-10 accuracy. Surprisingly, we therefore find that our algorithm actually performs better in predicting the pseudoknotted structures in the databases used than the non-pseudoknotted structures. The main results are the same for this dataset as for the full dataset plotted in Fig. 4: our algorithm outperforms all 14 algorithms tested against in predicting pseudoknotted structures, and performs on par with the other algorithms in predicting non-pseudoknotted structures, even though it uses orders of magnitude fewer entropic parameters than the other algorithms tested against.

**FIG. S4:**
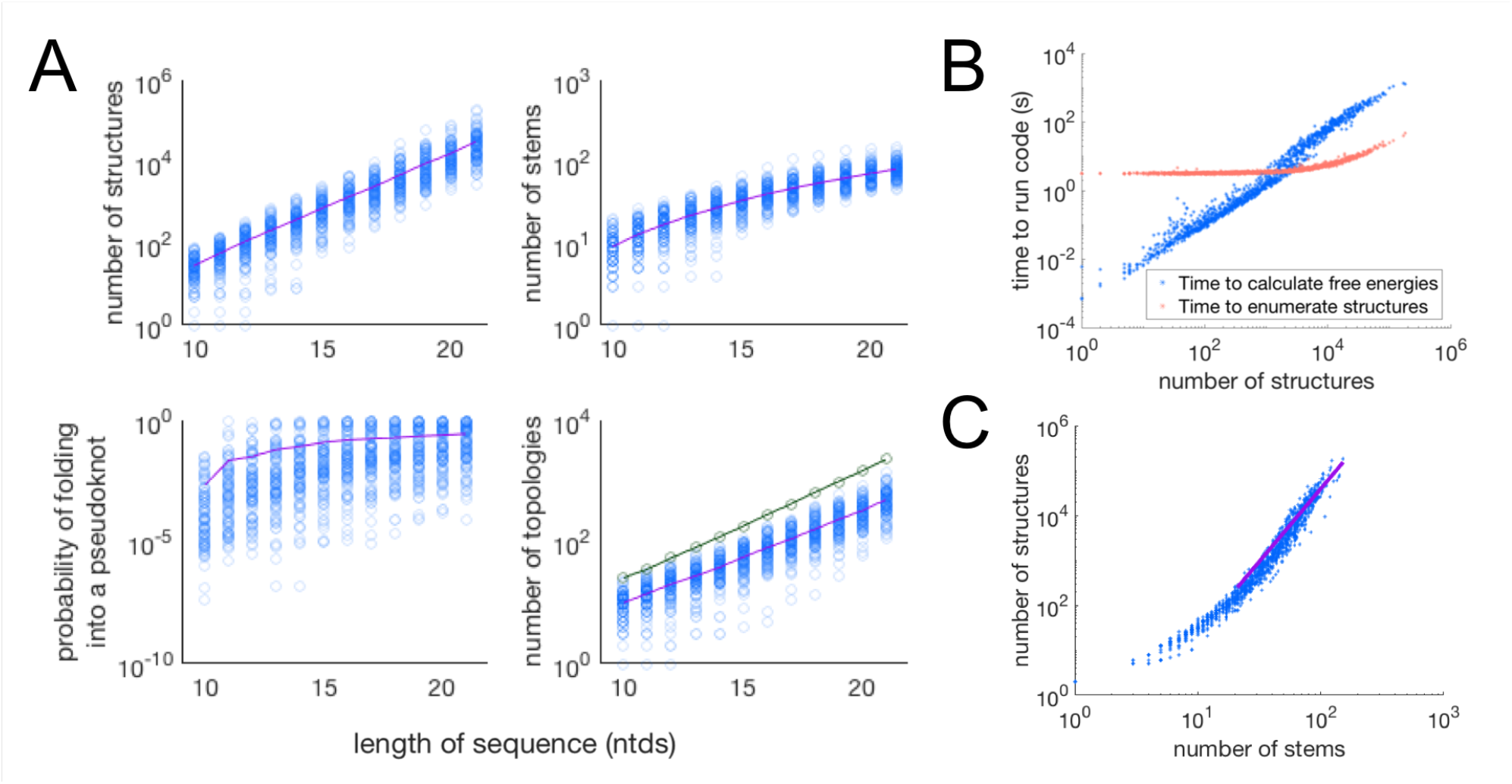
Scaling of the algorithm properties with length of sequence. We input 100 random sequences for each length between 10 and 21 nucleotides into the algorithm. (**A**) Various properties of the results are plotted as a function of the length of the sequence. Blue circles are datapoints for each of the 100 sequences in each column. Purple points show the mean. The number of secondary structures grows exponentially with the length of the sequence, as expected due to the NP-complete nature of the algorithm, though the number of possible stems grows sub-exponentially. The probability of forming a pseudoknot appears to plateau at around 10%. The number of topologies grows exponentially (we exclude topologies more complex than those shown in Fig. S2 and the structures leading to them). The green line shows the total number of different topologies over all 100 sequences of a given length. We disallowed parallel stems for this analysis. (**B**) The time the algorithm takes to calculate free energies grows approximately linearly with the number of possible secondary structures. The data is well-fit to a power law *y* = *ax^b^* with parameters *a* = (3.8 ± 0.3) * 10^−4^ and b = 1.27 ± 0.01. The time taken to enumerate all the structures is constant for short sequences (when few structures are enumerated and the algorithm’s overhead is the rate-limiting factor) and then grows as a power law. For sequences of any substantial length, the algorithm is rate-limited by the time it takes to compute free energies, rather than the time taken to enumerate structures. (**C**) For large numbers of stems, the number of possible secondary structures grows as a power law with the number of possible stems. This sub-exponential behavior is because some stems cannot coexist in the same structure (if they share any of the same nucleotides or if their coexistence leads to a topology more complex than those in Fig. S2). The purple line shows a fit to the equation *y* = *ax^b^* with *R*^2^ = 0.81. The best-fit values of *a* and b are found to be *a* = 0.0129 ± 0.0065 and *b* = 3.24 ± 0.11.

## Footnotes

1 More generally, we can define a probability 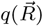 of a nucleotide at the origin being base paired with a nucleotide a vector 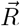 away. Then, *v_s_* is defined as 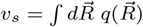 and *r_s_* is the value of 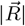 for which 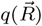 is non-negligible.

2 A more precise definition of *v_s_* might include a dependence on the closing base pairs of the hairpin loop; we expect that the penalties placed on specific closing base pairs and first mismatches in e.g. Refs. [64] and [25] play a similar role, though such penalties were not included here.

